# Regulation of the activity of an antimicrobial peptide by sterols or hopanoids: possible role in cell recognition

**DOI:** 10.1101/2020.06.23.166983

**Authors:** Dayane S. Alvares, Mariela R. Monti, João Ruggiero Neto, Natalia Wilke

## Abstract

It is now accepted that hopanoids act as sterol-surrogates in membranes of some sterol-lacking bacteria. Here we inquiry whether the hopanoid diplopterol (DP) could attenuate the activity of the antimicrobial peptide Polybia-MP1 (MP1) similarly to cholesterol (CHO). Survival of *P. aeruginosa* exposed to MP1 was lower for cells incubated with DP than those incubated with CHO, and the affinity and subsequent effect of the peptide on lipid bilayers were different in the presence of DP than in the presence of CHO. Membrane properties showed a non-monotonic behavior as the peptide adsorbed, penetrated, and translocated bilayers with DP suggesting a reorganization of MP1 during these processes. We conclude that MP1 selectivity is finely tuned by lipid composition, and propose the differential interaction and consequent effect promoted by the peptide in membranes with diplopterol as a promising starting point for targeting antimicrobial peptides to hopanoid-containing bacterial membranes.

## Introduction

Cell membranes are simultaneously compact and fluid. These properties, which are conserved in different species, have been proposed to be those that are compatible with life (Heimburg, 2007), and it is accepted that sterols are inducers of these particular mechanical conditions in different kingdoms. It has been shown in pioneer works and recent publications that hopanoids can be considered sterol-surrogates in sterol-lacking bacteria. Biophysical studies, using artificial membranes, have demonstrated that hopanoids decrease the membrane permeability and increase the membrane order without compromising their mechanical properties (Sáenz et al., 2012, 2015; Mangiarotti et al., 2019a). In addition, and in accordance with *in vitro* studies, hopanoids promote resistance to antibiotics, detergents, extreme pHs, high temperature, oxidation, and high osmolarity in bacteria (Sáenz, 2010; Belin et al., 2018; Welander et al., 2010; Malott et al., 2014; Schmerk et al., 2011; Silipo et al., 2014; Kulkarni et al., 2013).

Given the similarities between membranes with hopanoids and those with sterols, here we compared the activity of a known antimicrobial peptide (AMP), Polybia-MP1 (MP1, IDWKKLLDAAKQIL-NH_2_), against membranes with diplopterol (DP, a major hopanoid) or cholesterol (CHO). This peptide, extracted from the venom of the Brazilian wasp *Polybia paulista*, exhibits bactericidal activity against Gram-positive and Gram-negative bacteria being non-hemolytic and non-cytotoxic (Souza et al., 2005). The aim of this study is to get an insight into the reasons for this peptide behavior, deepening the understanding of the dependence of membrane composition on the MP1-membrane interaction. We inquired whether diplopterol protects membranes from peptide action as good as cholesterol, which hinders membrane disruption by MP1 (Dos Santos Cabrera et al., 2008, 2012). We showed using *in vivo* assays that the presence of DP prevents the action of MP1 to a lesser extent than that of CHO. This correlated with our results on the impact of DP on peptide-membrane affinity, MP1-lytic activity, and mechanical properties of artificial membranes, which showed important differences in membranes with DP compared to those with CHO. Our results indicate that the presence of hopanoids instead of CHO may be a critical factor for directing binding of this peptide to bacterial with higher affinity than to mammalian cells.

## Results and Discussion

### *P. aeruginosa* cells exposed to DP are more sensitive to MP1 than those exposed to CHO

A previous study using *P. aeruginosa* (a hopanoid-lacking bacteria) has shown that cells preincubated with CHO or DP, exhibit decreased permeability and sensitivity to cell wall biosynthesis inhibitors, carbenicillin, and imipenem (Mangiarotti et al., 2019a). This behavior correlates with the changes induced by the presence of CHO or DP in the properties of artificial lipidic membranes. In order to determine if DP and CHO provide an increased resistance to cells upon the action of MP1, the antimicrobial activity of this peptide against *P. aeruginosa* was evaluated by determining cell survival in cultures preincubated with CHO or DP. Cell survival to MP1 of untreated bacteria was similar to the control cultures previously exposed to the lipid solvent (Figures 1A and SI 1). In the presence of CHO, cells became insensitive to MP1 at 15 and 30 *µ*g/mL, being necessary to increase MP1 dose to 50 *µ*g/mL in order to observe cell death (data not shown). On the contrary, DP elicited a higher sensitivity to MP1 compared to CHO, as showed by the lower fraction of surviving cells. These data indicate that DP and CHO induce distinct MP1 sensitivities on bacteria. If this is related to different MP1 effects on membrane properties, MP1 should affect differently artificial membranes with DP or CHO.

**Figure 1.**
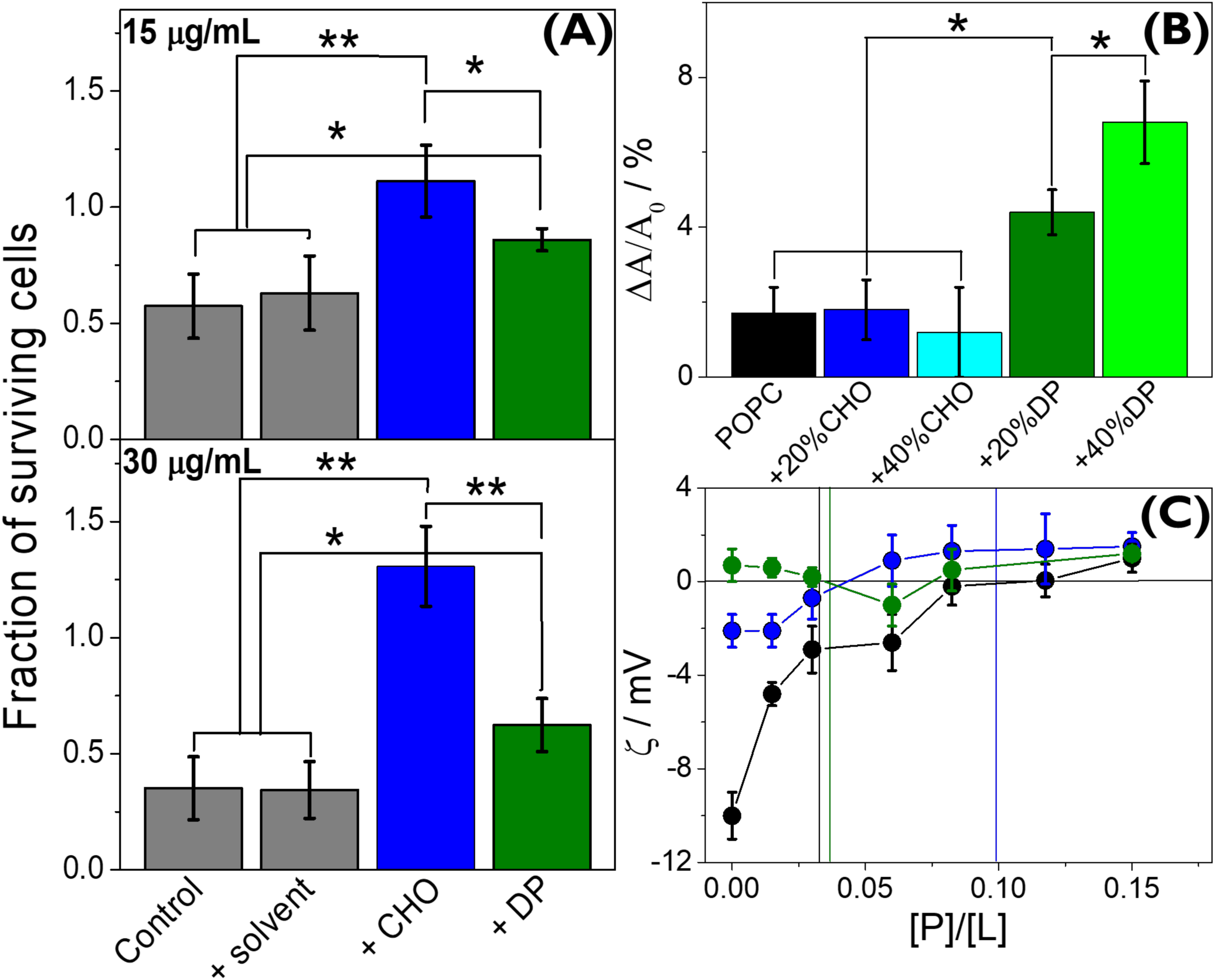
(A) Activity of MP1 against *P. aeruginosa* exposed to CHO or DP. Fraction of surviving cells without treatment (control) or incubated in the solvent, or a solution of CHO (blue) or DP (olive) after 3 hours of exposure to 15 *µ*g/mL or 30 *µ*g/mL of MP1. (B) Area change supported by lipid monolayers after peptide addition. Film composition: pure POPC (black), POPC + 20% (blue) or 40% (cyan) of CHO, or POPC + 20% (olive) or 40% (green) of DP. (C) Effect of MP1 on the zeta potential (*ζ*) of vesicles. *ζ*values for LUVs of pure POPC (black), POPC + 20% of CHO (blue) or 20% DP (olive) at increasing peptide/lipid ratio. The vertical lines represent the [P]/[L]_50_ values obtained from dye release experiments with CF loaded LUVs (Figure SI 3A). All data correspond to the average (± SD) of at least three independent experiments. * and ** indicate significant statistical differences determined by one-way ANOVA, Tukey’s test at p < 0.05 or p < 0.001, respectively.

### DP increases the perturbation induced by MP1 on lipid monolayers

We explored the process of insertion of MP1 into the external hemilayer of membranes using Langmuir monolayers. For 0.6 *µ*M peptide concentration, the maximum insertion pressure *π*_*MIP*_ (maximal value of surface pressure at which peptides insert into the film) was similar for all tested monolayers, although slightly higher for monolayers with 20% DP than those with the same amount of CHO (Figures SI 2A and 2B). Given the small differences found in *π*_*MIP*_, we decided to test films with higher amounts of CHO or DP at 30 mN/m, where the monolayer lipid compaction is comparable to that in bilayers (Demel et al., 1975; Marsh, 1996). Figure 1B shows the area change promoted in the lipid monolayer upon peptide penetration (see Alvares et al. (2018); Via et al. (2018) for details regarding calculation, and the changes in surface pressure in Figure SI 2C). Molecules that incorporate into monolayers have been shown to form interfacial structures with the lipids (Via et al., 2018; Vollhardt and Fainerman, 2000; Arouri et al., 2011) and to modify the film rheological properties differently depending on the host film (Alvares et al., 2016, 2017; Petit et al., 2009; Gidalevitz et al., 2003). Our results showed that, while increasing CHO levels did not result in changes on lipid-packing perturbation, the increase in DP levels promoted larger area changes suggesting that DP enhances the sensitivity of the membrane to this AMP. To test whether this is also the case for vesicles, we next investigated MP1-induced vesicle leakage.

### Dye release from liposomes is not affected by DP, while membrane electrostatics is affected in a complex fashion

The loss of vesicle content promoted by MP1 was assessed by recording the fluorescence of carboxyfluorescein (CF) released from large unilamellar vesicles (LUVs) (Figure SI 3A). MP1-induced leakage of POPC and POPC + 20% DP LUVs occurred without changes in the vesicle diameter, while those with 20% CHO formed larger aggregates at high peptide/lipid ratios ([P]/[L]) (Figure SI 3B).

The [P]/[L] value that induced 50% of CF release ([*P*]/[*L*]_50_) was 3-fold higher for membranes with 20% CHO than for LUVs of pure POPC (see vertical lines in Figure1C and Figure SI 3A). This has been previously related to a decrease in the probability of pore/defects formation by the AMP in CHO-containing membranes (Nir and Nieva, 2000; Lebarron and London, 2016). Opposite to the CHO-induced resistance, [*P*]/[*L*]_50_ for liposomes composed of POPC + 20% DP was similar to those of pure POPC. Monolayer experiments indicated that MP1 promoted the largest perturbations on DP-containing membranes, but this was not translated to differences in [*P*]/[*L*]_50_.

The addition of MP1 to POPC LUVs led to changes in the zeta potential (*ζ*), going from negative to positive values (Figure 1C). This was expected because, even though POPC is neutral, preferential binding of anions (Cl^−^ in our case) has been proposed, generating vesicles with negative *ζ* (Makino et al., 1991; Klasczyk et al., 2010; Manzini et al., 2014). Being a cationic peptide (net charge +2), MP1-adsorption neutralized the negative charge on the vesicles.

Despite *ζ* values at high [*P*]/[*L*] were similar in the three systems, the total changes were lower for LUVs with CHO and negligible for those with DP suggesting differences in the ionic cloud of the vesicles with the sterol or the hopanoid. The behavior of *ζ* for vesicles with DP was intriguing since a non-monotonically response was detected as [*P*]/[*L*] increased.

We will return to this point later. To get an insight into the peptide-bilayer affinity, the interaction of a fluorescently-labeled peptide was tested using giant unilamellar vesicles (GUVs).

### The luorescently-labeled MP1 distinguishes membranes with DP from those with CHO

A fluorescently-labeled MP1 (fMP1) was added to the GUVs in the observing chamber. Soluble peptides first adsorb to the membrane surface, penetrate the external hemilayer, and eventually move to the internal hemilayer, form pores and translocate to the GUV’s lumen. We followed the kinetics of these processes through the fluorescence of fMP1 at conditions comparable to the typical *in vivo* assays using AMPs.

Fluorescence of fMP1 was recorded during peptide binding at the rim of GUVs, and outside and inside them as shown in Figure SI 4. In Figure 2A, a POPC GUV is shown (other compositions are shown in Figures SI 5 and SI 6). The fluorescence outside a GUV quantifies the increase in fMP1 concentration close to the GUV due to peptide diffusion and the fluorescence in the interior measures the entry of the peptide into the GUV’s lumen. Rim’s fluorescence was normalized by the fluorescence outside each GUV to get an insight into the amount of accumulated peptide on the surface (*F*_*n*_). Considering a negligible change of the emission spectra of the fluorophore in different environments, values of *F*_*n*_ higher than one indicate that fMP1 concentration at the membrane is higher than outside the vesicle, thus peptide accumulates at the membrane.

**Figure 2.**
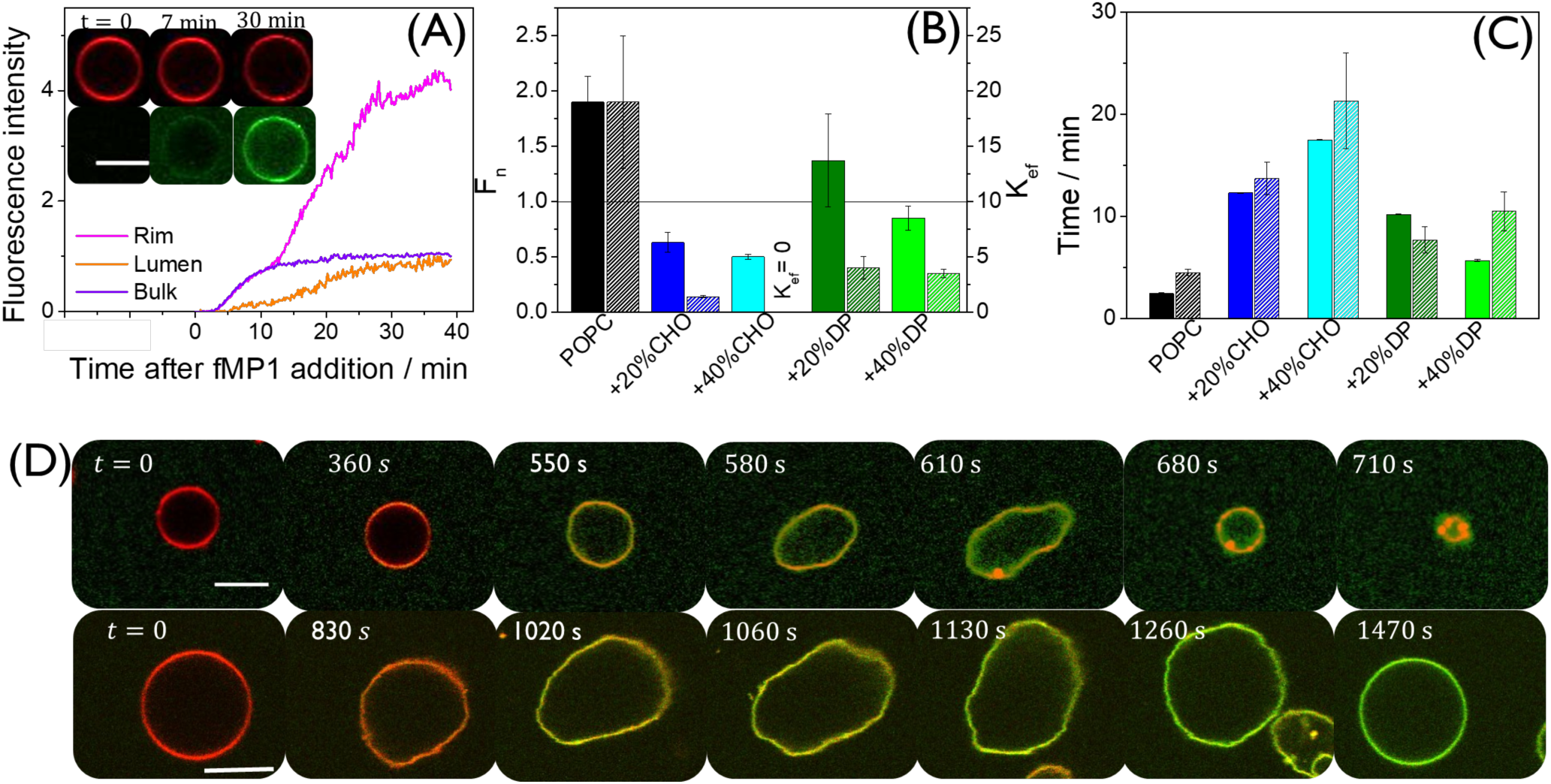
(A) Kinetic curves for the fluorescence intensity signal of MP1 labeled with FITC (fMP1) at the rim (magenta), at the lumen (orange) and outside a vesicle (violet) normalized by the final fluorescence outside the vesicle. GUV composition: POPC + 0.5 mol% of PE-Rhodamine in the presence of 0.6 *µ*M fMP1. Inset: confocal images at selected times after peptide addition. Upper sequence: red channel (PE-Rhodamine), lower sequence: green channel (fMP1). Scale bar: 10 *µ*m. For better visualization of the images, the brightness and contrast range was reduced from an original range of 0–255 to 0–132. (B) Final peptide fluorescence intensity at the membrane rim normalized by the intensity outside the GUVs (*F*_*n*_) (closed bars, left scale); and ratio between the constant for the rate of adsorption from outside and of desorption into the lumen of GUVs, K_*ef*_ (dashed bars, right scale). Data correspond to the average (± SD) of at least 30 (*F*_*n*_) or 15 (K_*ef*_) GUVs. (C) Times for achieving half the value of *F*_*n*_ at the vesicle’s rim (closed bars), and times at which peptide fluorescence is detected at the lumen of 50% of the vesicles (dashed bars). All data correspond to the average (± SD) of at least 45 GUVs. (D) Confocal images at selected times after addition of 6.0 *µ*M fMP1 to POPC vesicles (upper panels) or POPC + 20% DP vesicles (lower panels). The images correspond to the merge of the green (fMP1) and the red (PE-Rhodamine) channels. Brightness and contrast range: 0-150, scale bar: 10 *µ*m.

As shown in Figure 2B, *F*_*n*_ is the smallest for CHO-containing vesicles, in agreement with the lowest *A* values (Figure 1B). The peptide did not adsorb to some GUVs composed of 20% CHO and accumulated on the surface of others, followed by peptide entry into the lumen, thus showing a stochastic behavior (Figure SI 6). In the presence of DP, *F*_*n*_ values were higher than those for membranes with the same amount of CHO but smaller than for pure POPC membranes, thus, the presence of the hopanoid induced a decrease in peptide accumulation on the membrane. On the other hand, monolayer perturbation increased when DP was present in comparison to pure POPC films (Figure 1B). These opposing trends may be responsible for the lack of influence of DP on the peptide-induced leakage from LUVs.

The constant corresponding to the rate of peptide adsorption to the membrane from outside a vesicle (*k*_*a*_), and that for the rate of desorption from the membrane into vesicle lumen (*k*_*d*_) were estimated through the time evolution of the fluorescence intensities of the peptide at the GUV’s rim and outside the vesicle (see Material and Methods and Figure SI 7A). The ratio between *k*_*a*_ and *k*_*d*_ (*K*_*ef*_) gives an insight of the amount of peptide accumulated in the membrane, and shows a trend similar to *F*_*n*_ (Figure2B). In the case of pure POPC, *k*_*a*_ was higher than *k*_*d*_, indicating that the peptide remained in the membrane, probably forming peptide aggregates. In vesicles with 20% CHO, large values for both, *k*_*a*_ and *k*_*d*_ were obtained (Figure SI 7B), indicating that peptide adsorbed/desorbed fast, without remaining at the membrane. It is important to note that for this composition, the values of *k*_*a*_ and *k*_*d*_ were determined only for vesicles with appreciable peptide accumulation. Addition of 20% DP to POPC vesicles induced a decrease in both constants, while higher DP percentages increased their values, thus showing a non-monotonic behavior. For both DP concentrations, peptide accumulation in the membrane was higher than in membranes with CHO and lower than in pure POPC membranes.

The rate at which peptide entries into vesicles with DP was between that for pure POPC vesicles and those with CHO (Figure SI 8), indicating that despite MP1 induced similar permeability to the soluble CF species in membranes with or without DP (Figure SI 3A), peptide entry was affected by the presence of DP (Figure SI 8). Figure 2C depicts the times at which 50% of the GUVs had peptide in their lumen, together with the times at which the rim fluorescence intensity was 50% of *F*_*n*_. The similarity between the values indicates that peptide adsorption is highly correlated with peptide entry.

In summary, peptide accumulation inside GUVs derives from adsorption from the outside followed by slower desorption into the lumen. High peptide accumulation correlates with a high probability of peptide entry. For POPC and POPC + DP GUVs, we propose that the longer times of residence inside the membrane are related to the formation of peptide dimers/multimers, which led to development of pores/defects, and eventually, peptide entry and high membrane permeability.

No visible thermal shape fluctuation was detected for all GUV compositions before or after the addition of 0.6 *µ*M peptide (Figure SI 12). On the contrary, experiments with a 10-fold higher concentration of peptide revealed strong shape fluctuations (Figures 2D, SI 9, SI 10, and SI 11 and Movie SI 1. Furthermore, visible local lipid/peptide aggregates were detected, followed by membrane disintegration in GUVs composed of POPC and POPC + 20% CHO. In the latter, fluctuations started at larger times after peptide addition, and peptide-lipid aggregates followed by vesicle rupture was also observed at longer times after fluctuation started (Figure SI 12).

An interesting behavior was observed for membranes containing 20% DP in the presence of 6.0 *µ*M fMP1. The peptide first induced strong membrane fluctuations, and afterward, membranes returned to their initial state, without visual peptide/lipid aggregation or vesicle rupture (Figures 2D, SI 11, and SI 12 and Movie SI 1).

According to Meleard et al. (Méléard et al., 1997), from the factors characterizing membrane mechanical properties introduced by Helfrich (Helfrich, 1973), membrane thermal fluctuations in GUVs depend only on the bending elastic modulus *κ*_*c*_, and the spontaneous curvature *C*_0_. Thus, at high-dose, the peptide decreased *κ*_*c*_ and/or *C*_0_ for membranes with POPC and POPC containing 20% CHO leading to vesicle rupture. On the other hand, the same peptide levels induced first a decrease and then an increase in *κ*_*c*_ and/or *C*_0_, leading to stable vesicles that remain intact even at 6 *µ*M peptide. This intriguing non-linear effect was further studied at low concentration of the unlabeled peptide using a dynamic active method.

### The dynamic elasticity of membranes with different compositions is affected by MP1 in different manners

The effect of the peptide on membrane dynamic elasticity at low peptide dose (0.6 *µ*M) was investigated following the kinetic of retraction of membrane tethers (40 ± 5 *µ*m length) generated from single GUVs using optical tweezers (Movie SI-2). Values of [P]/[L] = 0.2-0.3 were used in these experiments, that is, higher than the value that induced 100% of CF release from LUVs.

The characteristic membrane relaxation time (*τ*) was determined by fitting curves such as that shown in Figure 3A with the equation 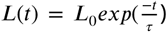 (see Material and Methods). The static force that oppose tether growth is 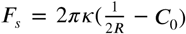 (Derenyi et al., 2007). At non-static situations, tether formation requires an additional force *F*_*d*_ which accounts for dissipation (Evans and Yeung, 1994). In tether retraction, *F*_*s*_ is the driving force while *F*_*d*_ oppose the motion. Thus, 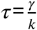, where *k* accounts for the elastic terms and depends on *κ*_*c*_ and *C*_0_, and *γ* is the dissipation coefficient. In our system, *γ* is formed by Stokes dissipation of the bead in the aqueous media, membrane shear viscosity (*η*_*s*_), and interlayer dissipation that depends on the coupling between hemilayers (*η*_*c*_) (Evans and Yeung, 1994; Hochmuth et al., 1996). Therefore, the rate of tether retraction decreases (and *τ* increases) if *κ*_*c*_ or *C*_0_ decreases, or if *η*_*s*_ or *η*_*c*_ increases.

**Figure 3.**
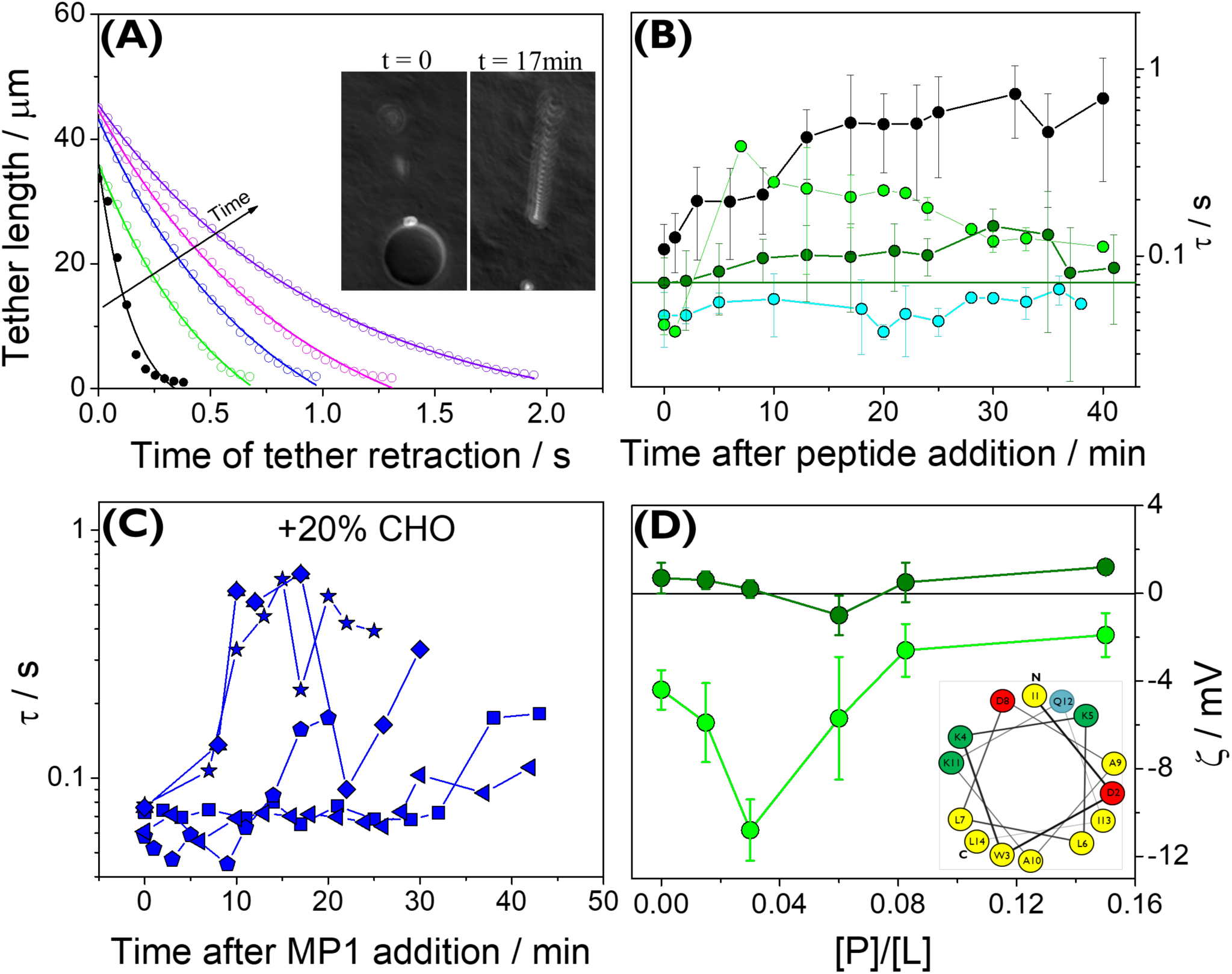
Membrane dynamic elasticity. (A) Retraction process of tethers pulled from a POPC GUV after 0, 10, 13, 17, and 25 min of peptide addition. The solid lines represent fits to the data using 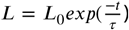, where *τ* is the characteristic time, 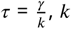, *k* is the elastic constant of the membrane and *γ* is the dissipation term. Inset: accumulated images obtained using DIC showing the *k*bead motion (24 frames/s) after the laser was switched off. Left: without peptide, right: after 17 min of MP1 addition, note the slower bead motion and the loss of vesicle contrast due to the increased permeability. (B and C) Characteristic times *τ* as a function of time after the addition of 0.6 *µ*M peptide to GUVs composed of (B) POPC (black), POPC + 20% DP (olive), POPC + 40% CHO (cyan) or DP (green), and (C) POPC + 20% CHO. Data in B correspond to the average (± SD) of at least 5 GUVs from at least two independent experiments. (D) *ζ*values for LUVs of POPC + 20% DP (olive) or 40% DP (green). Inset: wheel representation of MP1 in an *α*-helix configuration (green represents positively charged; red, negatively charged; blue, polar uncharged and yellow, hydrophobic residues, and N and C indicate the N-terminus and C-terminus of the peptide, respectively.)

The values of *τ* in the absence of peptide were higher for pure POPC in comparison to POPC-CHO membranes (see Figures 3B, C, and SI 14). This is expected since CHO induces an increase in *κ*_*c*_ (Henriksen et al., 2004; Dimova, 2014). DP promoted the same changes in *τ* as CHO, indicating a comparable effect of this lipid on the mechanical properties of POPC bilayers.

For POPC, POPC + 40% CHO and POPC + DP (both proportions), all analyzed GUVs showed the same trend in *τ*, and thus, the values of *τ* for different GUVs after peptide addition were averaged and plotted in Figure 3B. On the contrary, the behavior of GUVs composed of 20% CHO was stochastic, and thus, the *τ* values for individual vesicles are shown in Figure 3C. In the absence of peptide, *τ* did not change in time nor due to consecutive tether formation-retraction (Figure SI 13).

For POPC membranes, 0.6 *µ*M MP1 promoted a progressive increase in *τ*, requiring about 5 minutes for half the total change. This time was similar to those shown in Figure 2C indicating a correlation between peptide accumulation in the membrane, peptide entry and membrane softening. This increase in *τ*, together with the effect on thermal fluctuations promoted by 6.0 *µ*M fMP1 suggest that the peptide decreases the bilayer’s elasticity, as previously reported for other AMPs (Bouvrais et al., 2008; Shchelokovskyy et al., 2011; Dimova, 2014). This observation has previously been related to local membrane thinning (Ludtke et al., 1995; He et al., 1996), decrease in the lipid-lipid cohesion energy (Fa et al., 2007), peptide-peptide interactions(Bassereau et al., 2012; Tristram-Nagle et al., 2010; He et al., 1996), decoupling of the hemilayers (Shchelokovskyy et al., 2011), entropy effects (Bivas and Méléard, 2003), or inhomogeneous spontaneous curvature (Häckl et al., 1997; Bouvrais et al., 2008; Fa et al., 2007; Vitkova et al., 2006; Pan et al., 2009; Pabst et al., 2007; Dimova, 2014; Bassereau et al., 2012; Tristram-Nagle et al., 2010).

Unlike the behavior of POPC bilayers, no change in *τ* values was observed when GUVs with 40% CHO were exposed to the peptide, in agreement with the low effect of MP1 on this membrane composition. GUVs with 20% CHO appear to represent a composition close to a critical point, being more sensitive to uncontrolled parameters such as the precise vesicle composition or the surface tension. Interesting, the behavior in *τ* correlates with the stochastic adsorption and entry of fMP1 into this GUV composition (Figure SI 6)

Surprisingly, a non-monotonic behavior was observed in DP-containing POPC membranes. In 20% DP-containing GUV, a subtle increase in *τ* was first observed, followed by a decrease after 35 min of MP1 exposure, reaching a final value close to the initial one. This effect was more marked for membranes with 40% DP, in which MP1 induced a fast increase in *γ* (5-10 min after peptide addition), followed by a slow decrease. This behavior points to the presence of two opposing factors with different kinetics. Non-monotonic effects on membrane elasticity as the amount of peptide increases has been previously reported for Colistin in a three-component membrane. The behavior was explained considering dosedependent specific peptide-lipid interactions (Dupuy et al., 2018). In those experiments, vesicles were prepared with the peptide, and thus, reported data are at equilibrium conditions and bilayers are symmetric. Here, we analyzed the time-evolution of membrane mechanical response. Our membranes are two-components, and DP is not able to form stable vesicles when pure. Therefore, despite we cannot discard the above-mentioned explanation, we propose an alternative hypothesis.

We will now return to the *ζ* values shown in Figure 1C, where a dual behavior was observed. Figure 3D shows the *ζ* values for LUVs composed of POPC containing 40 mol% DP together with those for 20% DP (same as in Figure 3D). A decrease, followed by an increase in *ζ* was observed for vesicles with 40% DP, similar to those with 20% DP. The value of [*P*]/[*L*] where the minimum occurred was lower than for 20% DP, and the minimum was more marked. This particular behavior in *ζ*indicates that the trend in surface electrostatics changed as the amount of peptide in the membrane increased, suggesting a dose-dependent organization of the peptide at the membrane. Despite MP1 has a positive net charge, it contains two acidic residues that arrange at the surface as indicated by the wheel representation shown in Figure 3D. If the peptide adsorbs parallel to the interface, the hydrophilic part is expected to remain exposed (Zemel et al., 2005). The decrease in *ζ*may be a consequence of a higher exposition of the acidic residues than the cationic ones, or due to the presence of counter-ions.

Zemel et al. analyzed the membrane perturbation free energy during peptide-membrane interaction for weakly-charged peptides, considering interfacially-adsorbed monomeric and dimeric peptide species, and the multi-peptide transmembrane pore state. They found that in the transmembrane pore state, the lipid perturbation energy per peptide is smaller than in the adsorbed state, and proposed that the gain in conformational freedom of the lipid chains is a central driving force for pore formation (Zemel et al., 2005). They also found a weak lipid-mediated gain in membrane perturbation free energy upon dimerization of interfacially-adsorbed peptides.

Taking into account their results, we propose that at low [*P*]/[*L*] ratios, peptides adsorbed at the membrane in a diluted regime, exposing the acidic residues and turning the surface more negative. This would happen until a critical [*P*]/[*L*] value was reached, which depend on DP percentage. As the amount of peptide increased at the interface, the membrane perturbation free energy increased, and eventually peptide dimers or multimers would emerge. We showed previously for MP1 that the presence of peptide-peptide interactions was stabilized by inter-molecular salt-bridges involving acidic residues (Alvares et al., 2016). Therefore, cluster formation at the membrane would imply changes in the surface electrostatics that may give rise to the increase in *ζ*. The fact that the [*P*]/[*L*] ratio for the minimal value of *ζ*was double for 20% in comparison to 40% DP indicates that as DP increased, the effect of monomeric peptides on the membrane perturbation free energy increased, and lesser amount of peptide was needed for multimer formation.

Going back to the retraction experiments, despite these are kinetic assays and therefore not directly comparable to the *ζ*data, we propose that at short times, a low amount of peptide accumulated in the external hemilayer in a diluted regime. This first stage is followed by the formation of peptide dimers/multimers until pores/defects form, allowing vesicle leakage and the passage of the peptide to the vesicle lumen. The pores/defects may be formed by lipids and peptides (disordered toroidal pore (Li et al., 2015), or with one peptide at each hemilayer, since the thickness of the bilayers (∼0.4 nm) (Olsen et al., 2013) are twice the length of the peptide (∼0.2 nm) (Alvares et al., 2016).

The increase in *τ* occurred up to longer times for 20% than for 40% DP because in membranes with 20% DP, the amount of peptide required for the reorganization was higher. When peptides are in the outer hemilayer in a diluted regime, they may decrease *κ* by decreasing the lipid chain order (Zemel et al., 2008; Fa et al., 2007), increasing *η*_*s*_ (Smith-Dupont et al., 2010) or decreasing the term 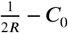 due to an increase in *C*_0_. This last effect is expected since at this stage membranes would be asymmetric. We discard effects due to *η*_*c*_ because a decrease, instead of an increase in hemilayer coupling is expected during this stage (Shchelokovskyy et al., 2011), which would lead to a decrease in *η*_*c*_ (and in *τ*).

When peptides reach a threshold concentration at the interface, they would form dimers/multimers, and eventually, pores. This would be accompanied by a decrease in *τ* that could be due to an increase in *C*_0_ since membrane asymmetry would decrease. Besides, a decrease in the effective *η*_*s*_ and an increase in the effective could occur in the presence of pores. In this sense, non-monotonic changes in *κ* have been proposed as the peptide moves inside the membrane (Zemel et al., 2008). Again, effects on the hemilayer coupling are discarded since, in the presence of pores, an increase and not a decrease in *η*_*c*_ is expected.

It is important to state that, despite the fact membrane dissipation terms cannot be neglected, the elastic response of the POPC/DP membranes likely commands the effect of the peptide on *τ*, since these data are in agreement with the changes in membrane elasticity detected by thermal fluctuations in GUVs composed of POPC + 20% DP in the presence of 6 *µ*M fMP1. The membrane-peptide association states proposed for POPC/DP membranes may also occur in pure POPC and POPC/CHO membranes. However, *ζ*and *τ* data indicate that membrane electrostatics and mechanical properties are affected in a different fashion in those membrane compositions.

## Conclusion

Our results show that the membrane composition is of paramount importance on the peptide-membrane affinity, as well as on the effect of the interaction on the properties of the host membrane, even in the case of neutral membranes. Lipid composition finely tunes the peptide-membrane interaction and also the consequences of this interaction on membrane destiny. Despite the similarities between the effect of DP and CHO on membrane properties, MP1 is able to exquisitely distinguish the presence of the hopanoid or the sterol, as well as their levels, being probably important for the selectivity of the peptide to hopanoid-containing bacteria from mammal cells. Although electrostatic interactions are likely to be important in the mechanism of the peptide-induced membrane break, the membrane mechanical properties are also key factors. The presence of anionic lipids may enhance the sensitivity of DP-containing membranes, and this should be investigated in turn.

## Methods

### Materials

Lipids: 1-palmitoyl-2-oleoyl-*sn*-glycero-3-phosphocholine (POPC), cholesterol (CHO), 1,2-dioleoyl-*sn*-glycero-3-phosphoethanolamine-N-(lissamine rhodamine B sulfonyl) (PE-Rhodamine), and 1,2-distearoyl-*sn*-glycero-3-phosphoethanolamine-N-[biotinyl(polyethylene glycol)-2000] (DSPE-PEG (2000)–Biotin) were purchased from Avanti Polar Lipids (Alabaster-Al-USA). Diplopterol (DP) was purchased from Chiron (Norway). Streptavidin-coated micro-beads (diameter of 3 *µ*m) were purchased from Bangs Laboratories Inc. (Fishers, IN, USA). MP1 peptide and MP1 with fluorescein isothiocyanate (fMP1) attached to the N-terminal were acquired from BioSynthesis (Lewisville-TX-USA) with RP-HPLC purity level > 98%. Chloroform HPLC grade was obtained from Merck (Darmstadt, Germany). Sodium chloride, sodium hydroxide, sucrose, HEPES, Triton X-100, carboxyfluorescein (CF), and avidin were from Sigma (St. Louis-MO-USA). Water used to prepare solutions was deionized with a resistivity of 18 MQcm, filtered with an Osmoion system (Apema, BA, Quilmes, Argentina).

### *In vivo* assay

Antimicrobial activity against *Pseudomonas aeruginosa*

The bacterial strain used in this study was *P. aeruginosa* PAO1. *P. aeruginosa* was cultured on Luria-Bertani (LB) agar plates from frozen stocks and one colony was grown in LB liquid medium at 37^*o*^C under shaking at 180 rpm overnight, up to an optical density of 1-1.2 at 600 nm. Cell suspensions were prepared in 0.15 M NaCl solutions to 2×10^4^ cells/mL. 14.4 *µ*L of pure chloroform, or 6 mM solution of Cholesterol or diplopterol dissolved in chloroform, was added to 120 *µ*L of the medium and left for 4h at 37^*o*^C for solvent evaporation. After that, 30 *µ*L of the media with lipids or solvent was added to 40 *µ*L of cell suspensions and incubated for 2h at 37^*o*^C without aeration. To evaluate susceptibility to MP1, cells were subsequently treated with 15 and 30 *µ*g/mL of MP1 and incubated for 3h at 37^*o*^C in a 96-multiwell plate, and plating into Luria-Bertani plates. Serial dilutions were plated on LB plates to quantify colony-forming units (CFU). The fraction of surviving cells was estimated from the ratio of colony-forming units (CFU) in the presence of peptide to that in its absence (control cells) as shown in Figure SI 1. For *in vivo* assay, the error bars of the fraction of cell surviving represent the deviations from the mean of the distinct experiments.

### Experiments using lipid monolayers

#### Insertion into lipid monolayers

The insertion of peptides into lipid films composed of pure POPC or the mixtures with DP or CHO were assessed in experiments at constant area using a home-made circular trough (volume 1 mL, surface area 7 cm^2^). An aliquot of concentrated MP1 solution was injected in the subphase above monolayers at an initial surface pressure (*π*_*i*_), through a hole in the wall of the trough. The final peptide concentration was 0.6 *µ*M. The adsorption kinetics of MP1 was followed by the increase in surface pressure (Δ*π*) as a function of time (Figure SI 2A). In order to obtain the maximum insertion pressure (MIP, surface pressure above which no more peptide molecules penetrate the lipid monolayer), Δ*π* induced by 0.6 *µ*M of MP1 as a function of different values of *π*_*i*_ was determined (Figure SI 2B). All experiments were carried out at 25 ^*o*^C in HEPES buffer (20 mM HEPES, 150 mM NaCl, pH 7.4). The standard deviations were obtained from at least three independent measurements.

### Experiments using large unillamelar vesicles (LUVs)

#### Preparation of LUVs

Multilamelar vesicles (MLVs) of pure POPC or containing 20 or 40% of CHO or DP were obtained from lipids dissolved in chloroform in round-bottom flasks. The solvent was dried under a stream of nitrogen and stored under vacuum for 3 hours to remove the organic solvent. Afterward, the lipid films were hydrated with a carboxyfluorescein (CF)-containing buffer and subjected to intense vortexing and several freeze-thaw cycles. LUVs were formed by extrusion of MLVs suspension, 15 times through a polycarbonate membrane of 0.1 µm pore size. The free dye was removed by exclusion chromatography using a Sephadex G-25M column (Amersham Pharmacia, Uppsala, Sweden) eluting the samples with HEPES buffer. To avoid osmotic pressure effects, osmolarity parity of CF and HEPES solutions was checked with an automatic micro-osmometer OM-806 (Vogel, Germany). The size distribution was confirmed by Dynamic Light Scattering (DLS) using a submicron particle sizer (Nicomp 380). The average diameter of the liposomes is shown in Figure SI 3B. In order to determine the actual amount of phospholipid after extrusion, the Bartlett’s method was used (Bartlett, 1958). Briefly, phospholipids were digested, and inorganic phosphorus was quantified colorimetrically. Absorbance was measured with a Shimadzu UV-visible Spectrophotometer (Biospec-mini, Chiyoda-ku, Tokyo, Japan) at *λ* = 830 nm.

#### Zeta Potential measurements

Zeta potential (*ζ*) of LUVs (40 *µ*M total lipid concentration) incubated with the peptide at increasing peptide-to-lipid molar ratio ([P]/[L]) were determined using a Z-sizer SZ-100-Z equipment (Horiba, Ltd., Kyoto, Japan). *ζ*of a vesicle suspension was calculated from electrophoretic mobility (*µ*) using Henry’s relation (Eq. 1):

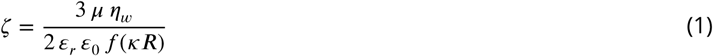

where *ε*_*r*_ and *ε*0 are the dielectric constant of water (78.5) and the vacuum permitivity, respectively, *η* _*ω*_ is the water viscosity, R is the average vesicle radius, is the inverse of Debye length and f(κR) in this case is 1.5 considering the Smoluchowsky approximation (Hunter, 1981). All experiments were performed at 25 °C. For these measurements, 15 mM NaCl was used as the aqueous media.

#### Dye release from CF-loaded LUVs

CF release from the vesicles was monitored detecting the dye emission at 520 nm (excited at 490 nm) on a Fluoromax-P spectrofluorometer (Horiba Jobin-Yvon). The maximum fluorescence intensity (100%) was determined by adding 10% Triton X-100 solution into a cuvette with the liposomes (complete membrane solubilization). The percentage of dye release induced by MP1 was calculated at regular time intervals using the Eq. 2 :

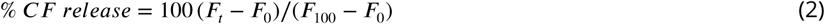

where F_*t*_ is the observed fluorescence intensity, F_0_ and F_100_ corresponds to the fluorescence intensities in the absence of peptides and for 100% leakage, respectively (Weinstein et al., 1981). The experiments were performed at 25 ^*o*^C. Vesicles prepared in the day were used in each experiment. Standard deviations were obtained from three independent experiments.

### Experiments using giant unillamelar vesicles (GUVs)

#### Preparation of GUVs

GUVs were prepared by the electroformation technique as described by Angelova and Dimitrov (Angelova and Dimitrov, 1986) using a home-made wave generator and a chamber with stainless steel electrodes (Bellon et al., 2018). Briefly, 7 *µ*L of a 0.5 mM lipid solution (in chloroform) were spread on two stainless steel electrodes, and were left under vacuum for at least 3h to remove all traces of the organic solvent. The mixtures contained the desired lipid composition plus 0.5% (molar ratio) PE-Rhodamine and 0.0001% (molar ratio) DSPE-PEG (2000)–Biotin. The lipid films were hydrated by filling a home-made chamber of acrylic containing the electrodes with a 0.3 M sucrose solution. The electrodes were connected to a wave generator, and a sinusoidal tension of 1-2 V amplitude and 10 Hz frequency was applied for 1-3 h at 25 ^*o*^C for POPC vesicles, and 60 ^*o*^C for vesicles with CHO or DP. GUVs were then harvested and added into a chamber containing an iso-osmolar HEPES buffer for further experiments. To avoid osmotic pressure effects, osmolarity parity of inside and outside solutions was checked with an automatic micro-osmometer OM-806 (Vogel, Germany).

#### Assay for peptide affinity to GUV

GUVs were directly observed under an inverted confocal laser-scanning microscope (Olympus Fluorview FV1000, Tokyo, Japan). Briefly, the GUVs were suspended in 250 *µ*L of HEPES buffer in an 8-well Lab-Tek Chamber. Then, confocal timeseries images taken with an oil immersion 60X objective (NA: 1.4, Olympus), were sequentially acquired with two lasers (Argon - 488 and Helium-Neon - 543 nm) before and after addition of fMP1 with the help of a micropipette to yield the desired peptide concentration. Peptide surface excess on the bilayers was estimated from the fluorescence intensity at the GUV’s rim in the green channel. This intensity was determined as a function of time by drawing a line at the vesicle contour (Figure SI 4). This intensity was normalized by the bulk intensity outside the GUV. To determine peptide penetration into the vesicle, the fluorescence intensity at the GUV interior was measured using a small circle in the center of the GUV as exemplified in Figure SI 4. Figures SI 5 and SI 6 show the signal quantification for all lipid composition, except for POPC which can be found in the manuscript. The fluorescence intensities were measured using the NIH free ImageJ/FIJI software.

#### Kinetic of peptide adsorption and entry into the vesicle lumen

The peptide added in the aqueous solution first diffuses to the vesicle, and then adsorbs at the membrane surface. Eventually, it penetrates into the outer and the inner leaflet, forms dimers or multimers, and pores/defects, and desorbs into the aqueous solution in the interior of the vesicle. In our experiments, the different peptide species in the membrane (dimers, multimers, pores) cannot be distinguished, and only the total peptide concentration in the membrane (*C*_*M*_) can be ascertained through the fluorescence intensity at the membrane rim. We will consider that initially, the reaction occurs in a unidirectional fashion: peptides adsorb (without desorption), accumulate in the membrane forming dimers, multimers or pores, and thereafter, the peptide aggregates disassemble and desorb (without readsorption) mainly in the vesicle lumen, where their concentration (*C*_*i*_) is initially zero. In other words, initially (for low peptide concentration in the vesicle lumen) we can neglect the backward reactions, and only consider adsorption of peptides from outside the vesicle to the external hemilayer and desorption from the internal hemilayer to the vesicle lumen. At these conditions, the following holds:

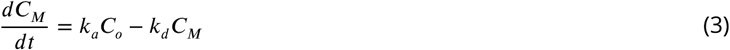

Here, *C*_*M*_ include all the possible peptide species inside the membrane, *C*_*o*_ is the peptide concentration outside the GUV, and *k*_*a*_ and *k*_*d*_ are the rate constants for adsorption from outside the vesicle and desorption into the vesicle interior, respectively. Since the local peptide concentration is proportional to the local fluorescence intensity (neglecting changes in the quantum yield of FITC with the environment), we can fit the data from fluorescence intensity at intermediate times (before *C*_*i*_ reached a constant value, and after *C*_*o*_ remained constant to avoid the diffusional regime) using Eq. 4:

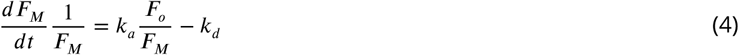

Here, *F*_*M*_ and *F*_*o*_ are the fluorescence intensities of the peptide at the vesicle rim and outside the vesicle, respectively. Using this equation, the kinetic of peptide adsorption and entry was fitted for experiments with 0.6 *µ*M fMP1 (see a representative plot in Figure SI 7A). Higher peptide concentrations were not used due to peptide-lipid aggregation, membrane fluctuations and vesicle rupture. The data for POPC + 40% CHO did not allow fitting due to the low fluorescence signal at the vesicle rim. From the obtained values, the ratio between the rate constants was determined 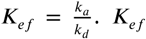 is a measure of the peptide ability to accumulate at the membrane.

#### Peptide entry to the GUV’s lumen

Peptides accumulate in the membrane, and eventually enter into the vesicle interior. We determined the amount of GUVs that had fMP1 in their interior at different times after peptide addition, and calculated the fraction of GUVs in which peptide had entered (*f*_*e*_) at each time (Figure SI 8). The analysis was performed using at least 15 GUVs in three independent experiments, Figure SI 8 shows the average of the three experiments ± SD.

#### Shape fluctuation analysis

Images were collected using two channels: red, to visualize the membrane marker (PE-Rhodamine), and green for the FITC-labeled peptide. Figures SI 9, SI 10, and SI 11 show a representative GUV sequence for POPC, POPC + 20% CHO, and POPC + 20% DP, respectively. Membrane shape was analyzed using NIH ImageJ/FIJI software. We determined the aspect ratio (AR, the ratio between the lengths of the major and minor axis) as a function of time, see Figure SI 13.

#### Optical Tweezer Setup

The used optical trap equipment has been described previously (Crosio et al., 2019). Briefly, a 3W power ND:YVO4 laser (Spectra-Physics, Santa Clara, CA, USA) was focused through a 100x objective (water immersion, NA = 1, Zeiss). The tweezers were mounted in an Axiovert 200 inverted microscope (Zeiss) equipped with a motorized stage (Beijing Winner Optical Ins. CO., LTD, Langfang, Hebei, China), and the experiments were registered with a fast CCD video camera (Ixon EM+ model DU-897, Andor Technology, Abingdon, Oxfordshire, England).

#### Membrane tether retraction assays

The experiments were performed in a Teflon homemade chamber with a coverslip (Marienfeld Superior, Germany) as optical window. The coverslip was treated with an avidin solution (0.5 mg/mL for 30 min at 10 ^*o*^C). The chamber was placed on the motorized stage of the microscope. A 3.0 *µ*m streptavidin-coated bead solution was added to the chamber containing HEPES buffer. After incubation of 10 min, a GUV solution was added to the chamber and left to attach to the avidin-coated glass due to avidin-biotin interactions. A bead was trapped with the optical tweezers and was brought into contact with the surface of a proximal GUV. Bead-membrane adhesion was achieved by means of streptavidin-biotin interactions. Once the bead adhered to the membrane, a membrane tether was pulled at a controlled speed of 15 *µ*m/s up to a length of 40 *µ*m. At t=0 the laser was switched off and tether retraction was registered by contrast microscopy at a rate of 24 frames/s. Movie S2 shows an example. A representative image sequence is shown in Figure SI 13.

#### Nanotube retraction analysis

Tether retraction was followed through the bead motion. The equation of motion of the bead can be obtained after applying Newton’s second law:

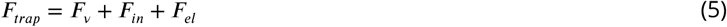

*F*_*γ*_ is the viscous force that opposes motion, and includes the dissipation of the bead in the aqueous media and membrane dissipation. Bead dissipation can be computed through the Stokes law: 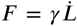, where *γ* = 6*πηR*_*b*_ = 28*nNs*/*m* (*R*_*b*_ is the radius of the bead). Membrane dissipation has the contribution of shear (*η*_*s*_) and of the friction between leaflets (*η*_*c*_), the sum of *η*_*s*_ and *η*_*c*_ is in the range of 1-100 nN s/m Evans and Yeung (1994); Hochmuth et al. (1996). *F*_*in*_ is the inertial term 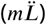, which can be neglected for low Reynolds numbers Marco Capitanio (2017), and *F*_*el*_ = *kL* corresponds to membrane elasticity, and is the driving force for retraction. When the trap is turned off, *F*_*γ*_ = –*F*_*el*_, leading to 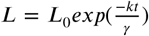, where *L*_0_ is the initial tether length (40 *µ*m). Since we cannot split the contribution of membrane dissipation from membrane elasticity, the characteristic time 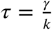 is obtained from the fitting process. The used fitting equation therefore is Eq. 6:

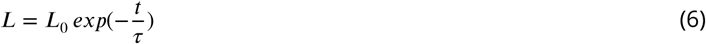

## Acknowledgments

The authors acknowledge financial support from UNESP, CAPES, CNPq, SECyT-UNC, FONCYT (PICT 2015-0662) and from São Paulo Research Foundation (FAPESP #2015/25619-9). D.S.A has a Post-doctorate fellowship grant #2015/25620-7 and BEPE-grant #2018/14215-2. J.R.N. is a researcher of CNPq. M.R.M and N.W. are researchers of CONICET-Argentina. We thank Dr. Mas and Dr. Sampedro for technical assistance at CEMINCO (Argentina).

## Supporting Information

### Movies

**Movie SI 1**. Time evolution of vesicles showing the shape fluctuations induced by 6 *µ*M MP1. GUVs are composed of pure POPC (left), POPC + 20% CHO (middle) or POPC + 20% DP (right). GUVs were observed for a total time of 350s (POPC), 480s (POPC + 20% CHO) or 650s (POPC + 20% DP). Scale bar: 10 *µ*m.

**Movie SI 2**. Tether retraction after being pulled from a single biotinylated-GUV immobilized onto an avidin-coated glass slide. GUV is composed of pure POPC, and the movie was registered before peptide addition (left) or after 13 min of the addition of 0.6 *µ*M MP1 (right). The video was captured during 0.3 and 0.6 s in the absence and the presence of peptide, respectively. Scale bar: 10 *µ*m.

### Figures

**Figure SI 1.**
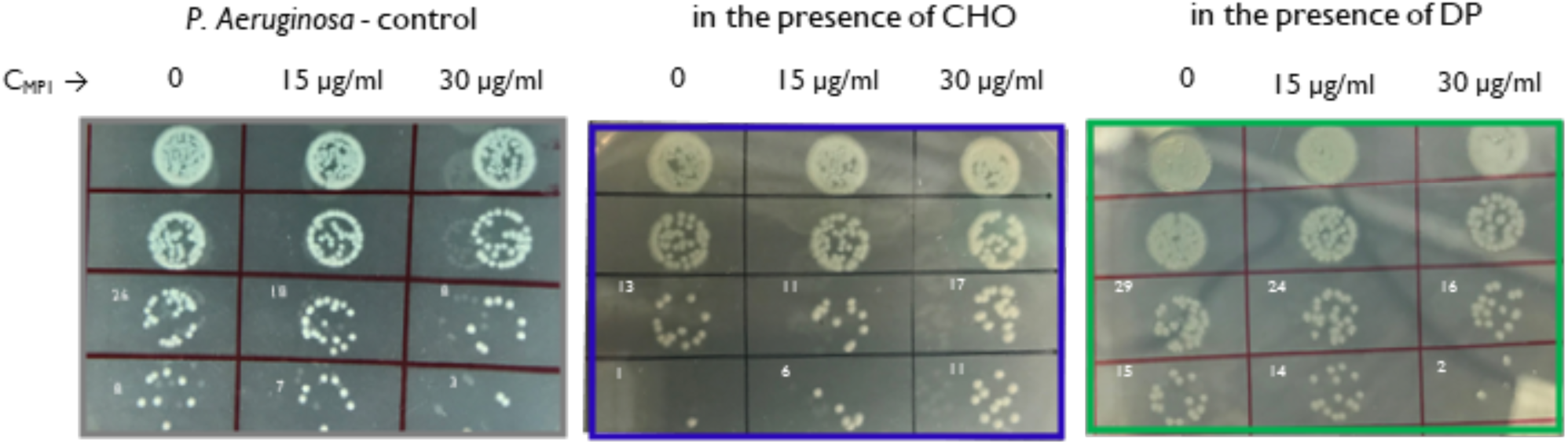
Representative photos showing the growth of colonies on agar plates. Each line represents serial dilutions, the first column represents bacterial strain in the absence of peptide, in the second and third columns the inocula were treated with 15 and 30 *µ*g/mL of MP1, respectively.

**Figure SI 2.**
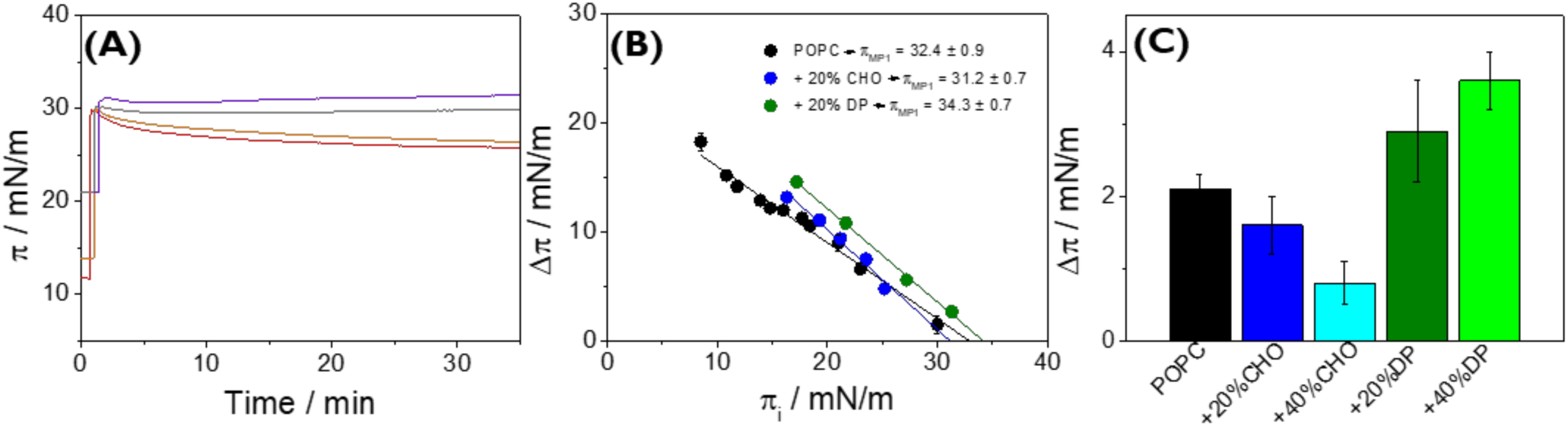
Insertion peptide’s ability into monolayers. (A) Insertion kinetics of the peptide into POPC monolayers at different initial surface pressures (*π*_*i*_). (B) Total change in the surface pressure (Δ*π*) after injection of 0.6 *µ*M MP1 in the subphase as a function of *π*_*i*_. Monolayers are composed of pure POPC (black), POPC + 20% CHO (blue) or POPC + 20% DP (olive). The continuous lines represent linear regressions. (C) Final change of Δ*π* after injection of 0.6 *µ*M MP1 beneath lipid monolayers at 30 (±0.5) mN/m. Monolayers are composed of pure POPC (black), POPC + 20% CHO (blue), POPC + 40% CHO (cyan), POPC + 20% DP (olive), or POPC + 40% DP (green). The subphase was HEPES buffer (20 mM HEPES, 150 mM NaCl) at pH 7.4. All data correspond to the average (± SD) of three independent experiments.

**Figure SI 3.**
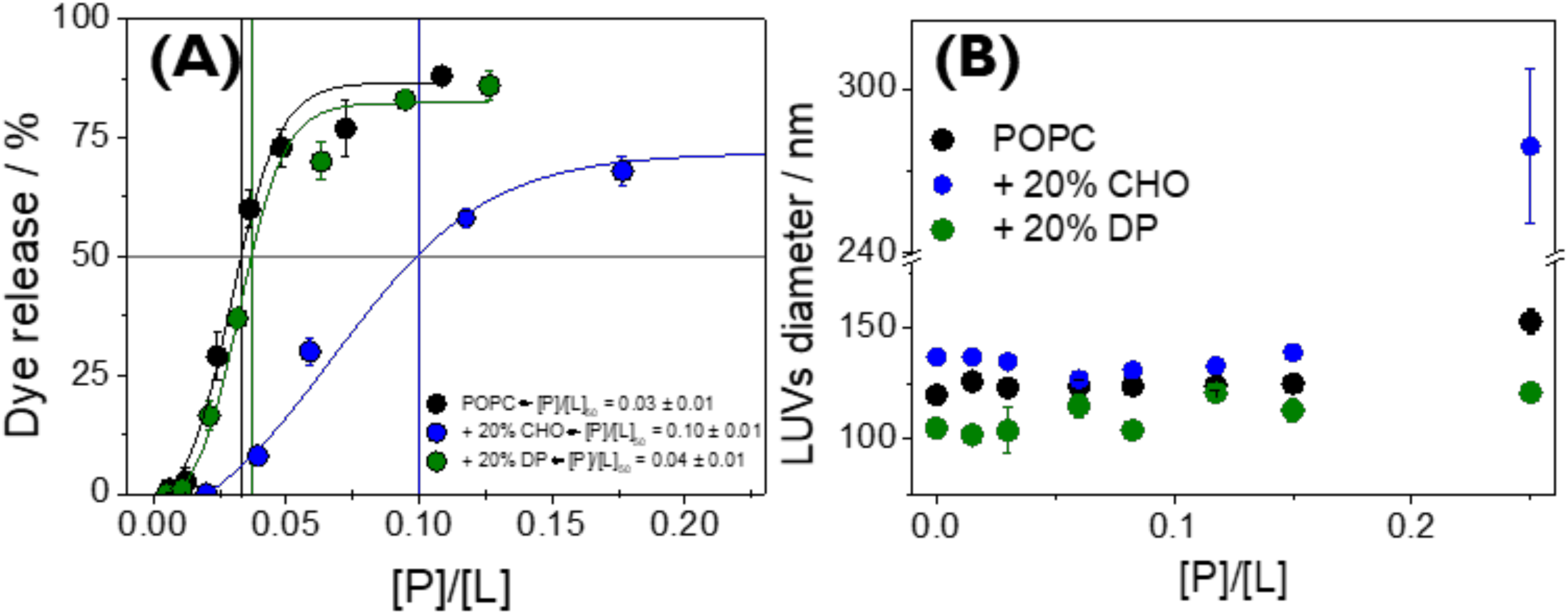
(A) Peptide’s lytic activity. Florescence of the released carboxyfluorescein (CF) from LUVs composed of pure POPC (black), POPC + 20% CHO (blue), or POPC + 20% DP (olive). The vertical lines represent the [P]/[L]_50_ values. (B) Changes in LUV’s average diameter as a function of peptide/lipid molar ratio. All data correspond to the average (± SD) of three independent experiments.

**Figure SI 4.**
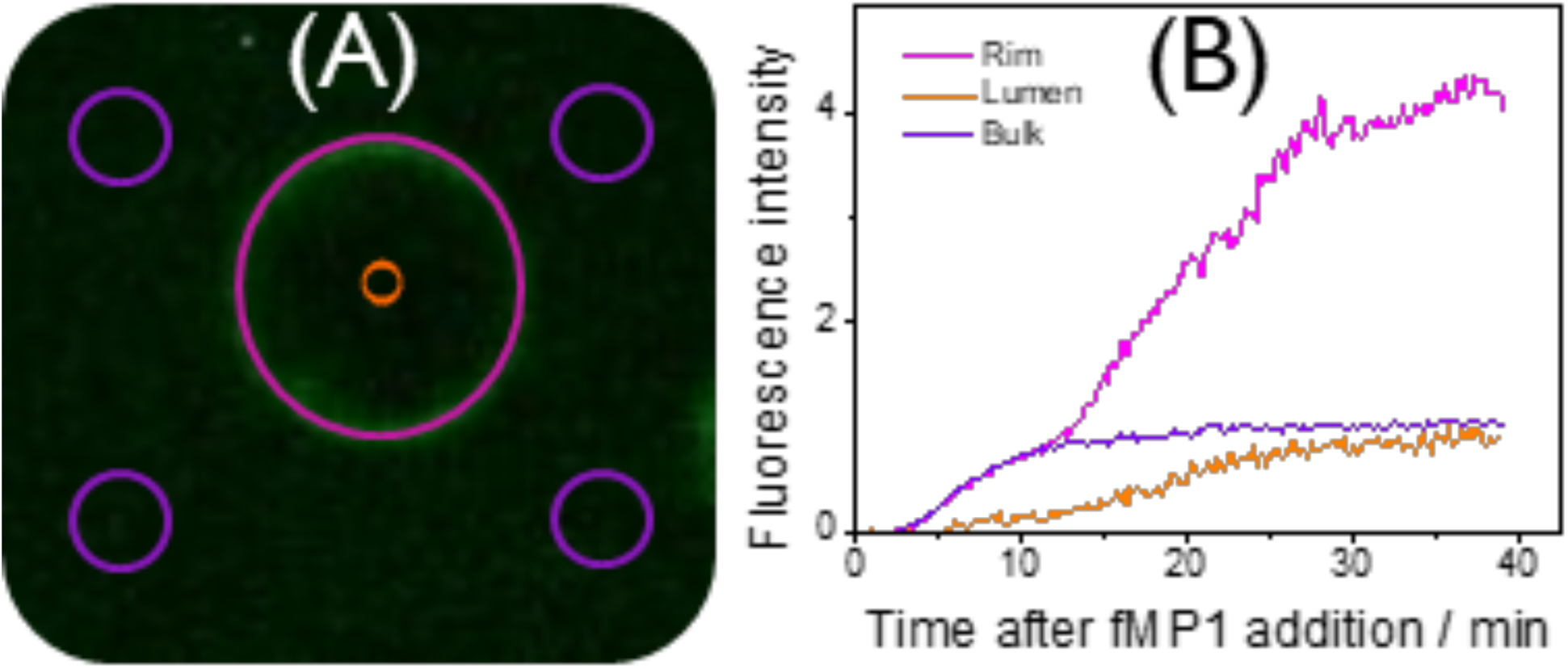
Measurement of peptide’s fluorescent intensity. (A) A GUV composed of POPC 20 min after the addition of 0.6 *µ*M of FITC-labeled MP1 (green signal). Fluorescence signal inside the violet circles depends on the peptide concentration in the bulk, outside the vesicle. The fluorescence at the magenta line gives an estimate of peptide accumulation at the membrane and the fluorescence inside the orange circle indicates the amount of peptide in the vesicle lumen. (B) Quantification of the fluorescence as shown in A gives curves of the time course of peptide intensities outside the vesicle (bulk), at the vesicle’s rim, and inside the vesicle (lumen).

**Figure SI 5.**
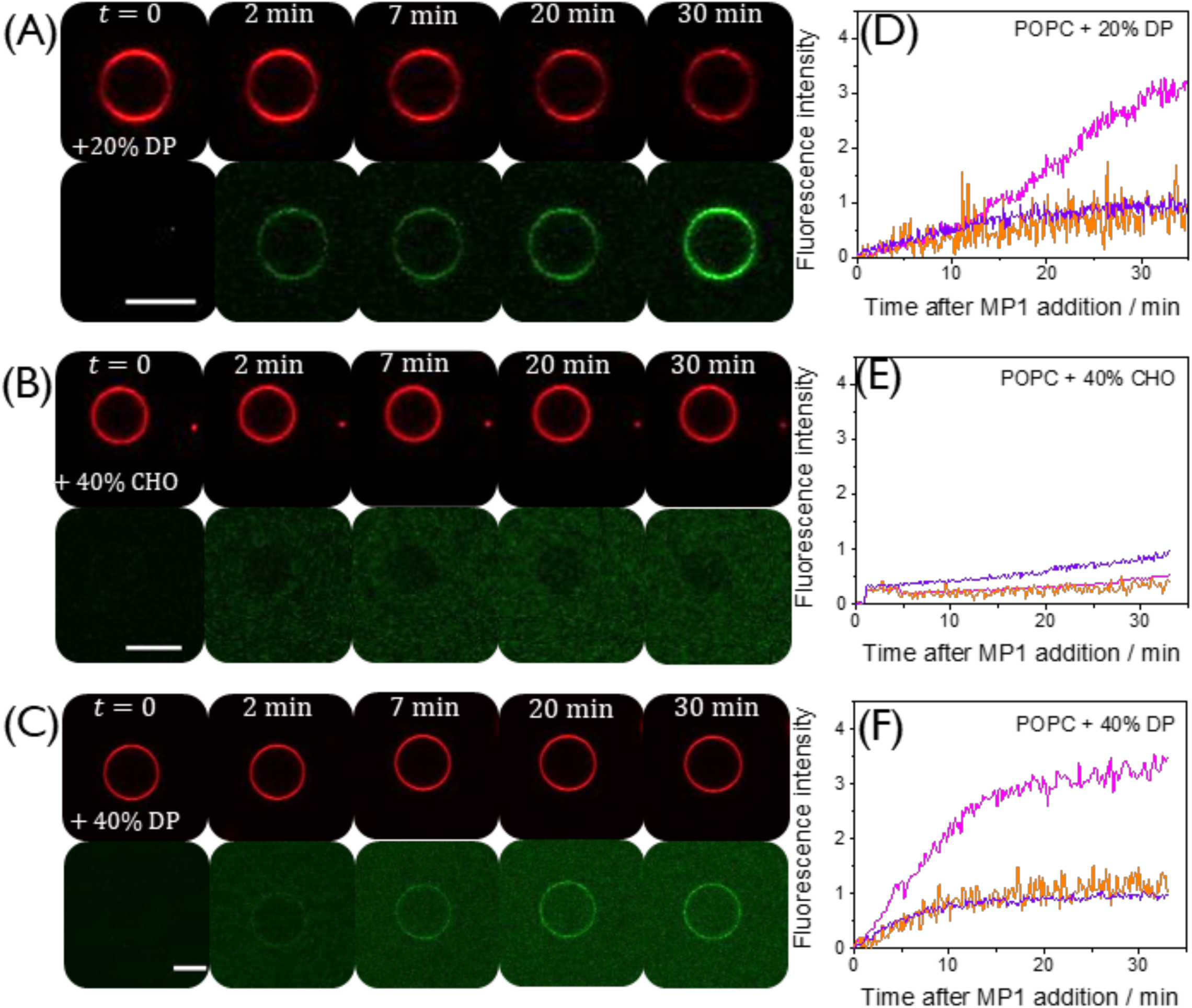
Confocal images of GUVs after fMP1 addition. (A-C) Representative confocal fluorescence images of a GUV composed of POPC + CHO or DP doped with 0.5 mol% of PE-Rhodamine (red signal). The indicated times correspond to times after addition of 0.6 *µ*M of fMP1 (green channel). Scale bar: 10 *µ*m. For better visualization of the images, the brightness and contrast range was reduced from an original range of 0–255 to 0–150, and the second image sequence in (B) was adjusted to 0-100. (D-F) Kinetic curves showing the fluorescence intensity of fMP1 at the vesicle’s rim (magenta), in the vesicle’s lumen (orange) or in the bulk solution (violet). Fluorescence intensities were normalized by the final fluorescence signal at the bulk after peptide addition.

**Figure SI 6.**
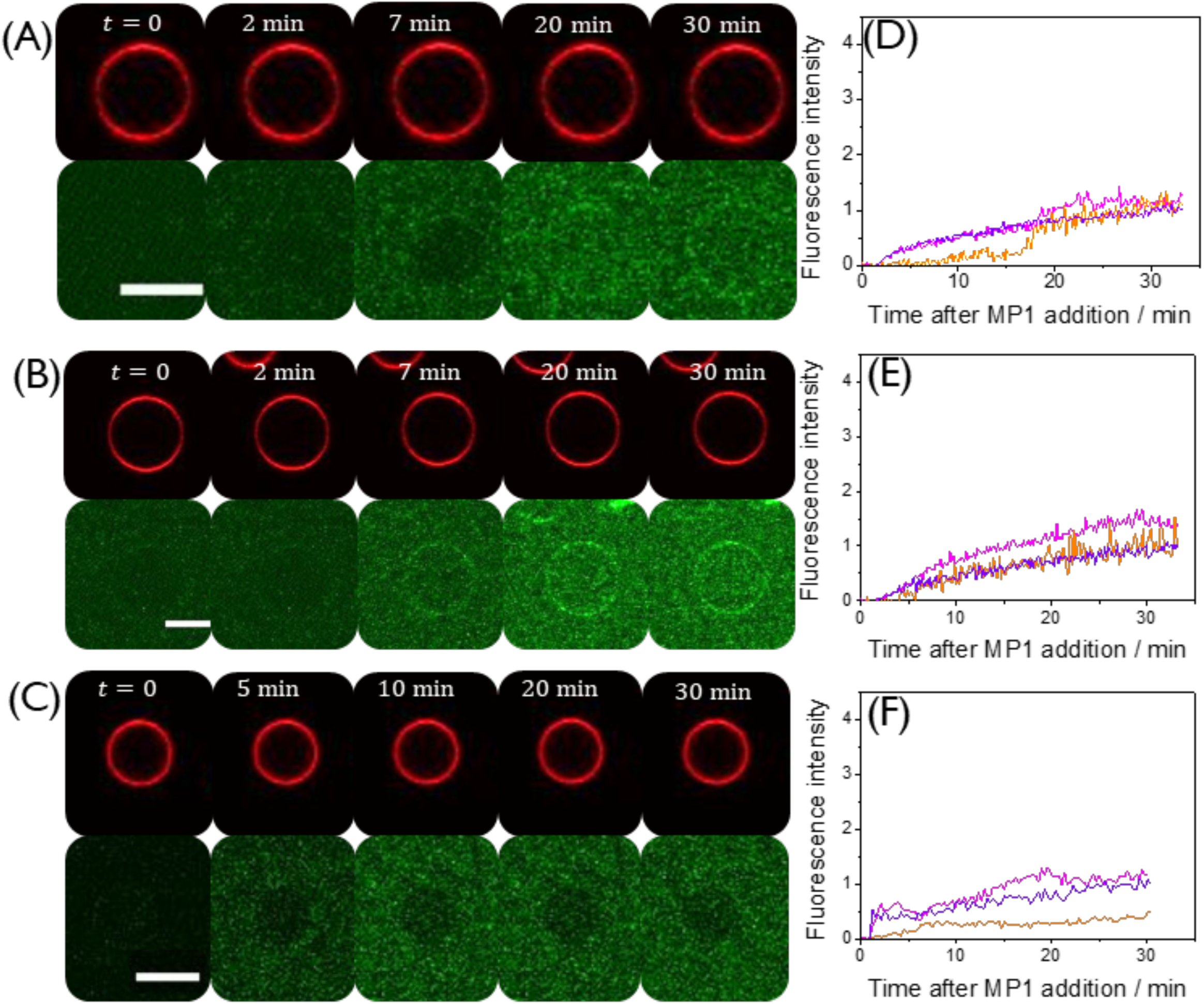
Confocal images of GUVs composed of POPC + 20% CHO after fMP1 addition. (A-C) Representative confocal fluorescence images of different GUVs doped with 0.5 mol% of PE-Rhodamine (upper panels – red signal) after addition of 0.6 *µ*M MP1 labeled with FITC (lower panels – green signal). Times after peptide addition are indicated. Scale bar: 10 *µ*m. For better visualization of the images, the brightness and contrast range was reduced from an original range of 0–255 to 0–150 (red signal) or 0-100 (green signal). (D-F) Kinetic curves for the fluorescence intensity of fMP1 at the vesicle’s rim (magenta), in the GUV’s lumen (orange), or in the bulk solution (violet). Fluorescence intensities were normalized by the final fluorescence signal at the bulk after peptide addition.

**Figure SI 7.**
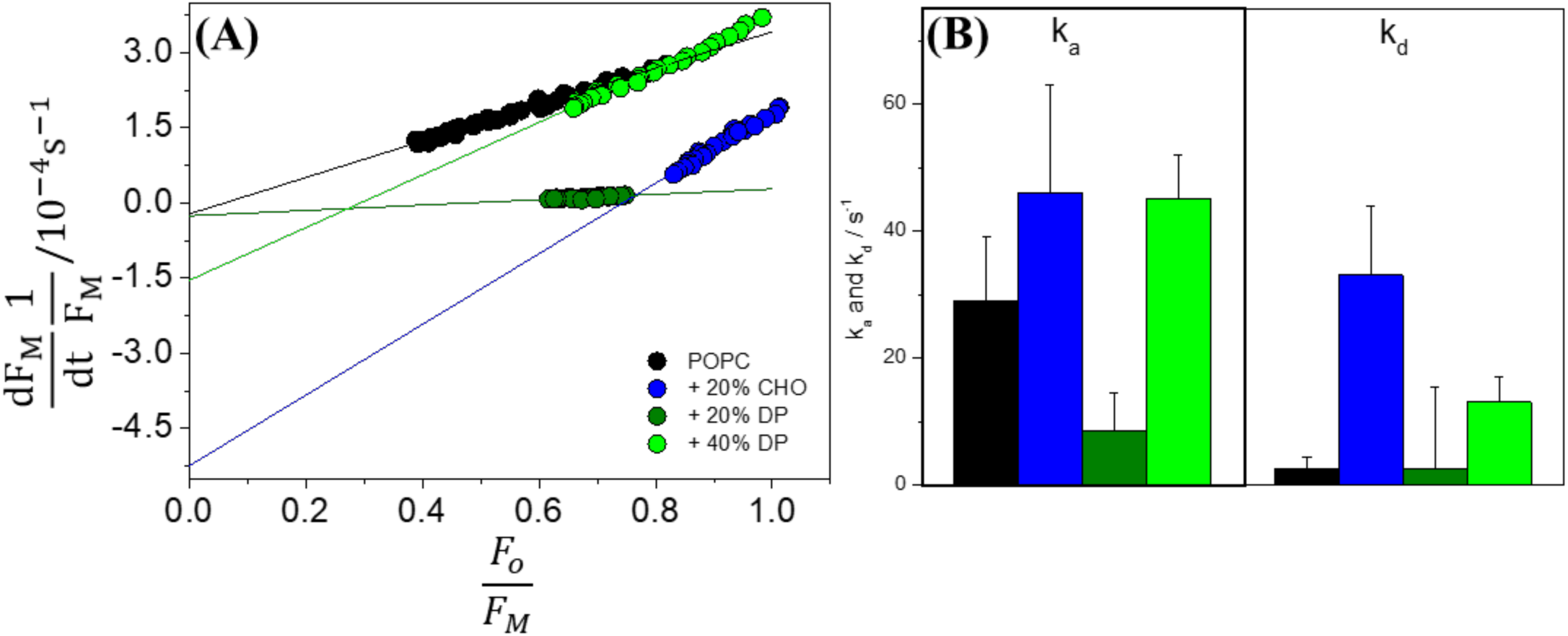
Kinetic analysis of peptide adsorption and entry. (A) Representative data from the fluorescence intensity of fMP1 for an experiment using a single GUV of the indicated composition, and the corresponding fit using Eq. 4. (B) Average rate constant for peptide adsorption from outside the vesicle (*k*_*a*_) or desorption to the vesicle’s lumen (*k*_*d*_) for GUVs composed of POPC (black), POPC + 20% CHO (blue), POPC + 20% DP (olive), or POPC + 40% DP (green). *k*_*a*_ and *k*_*d*_ values correspond to the average (± SD) of the data from 10 single GUVs, such as those shown in (A), obtained from two independent experiments.

**Figure SI 8.**
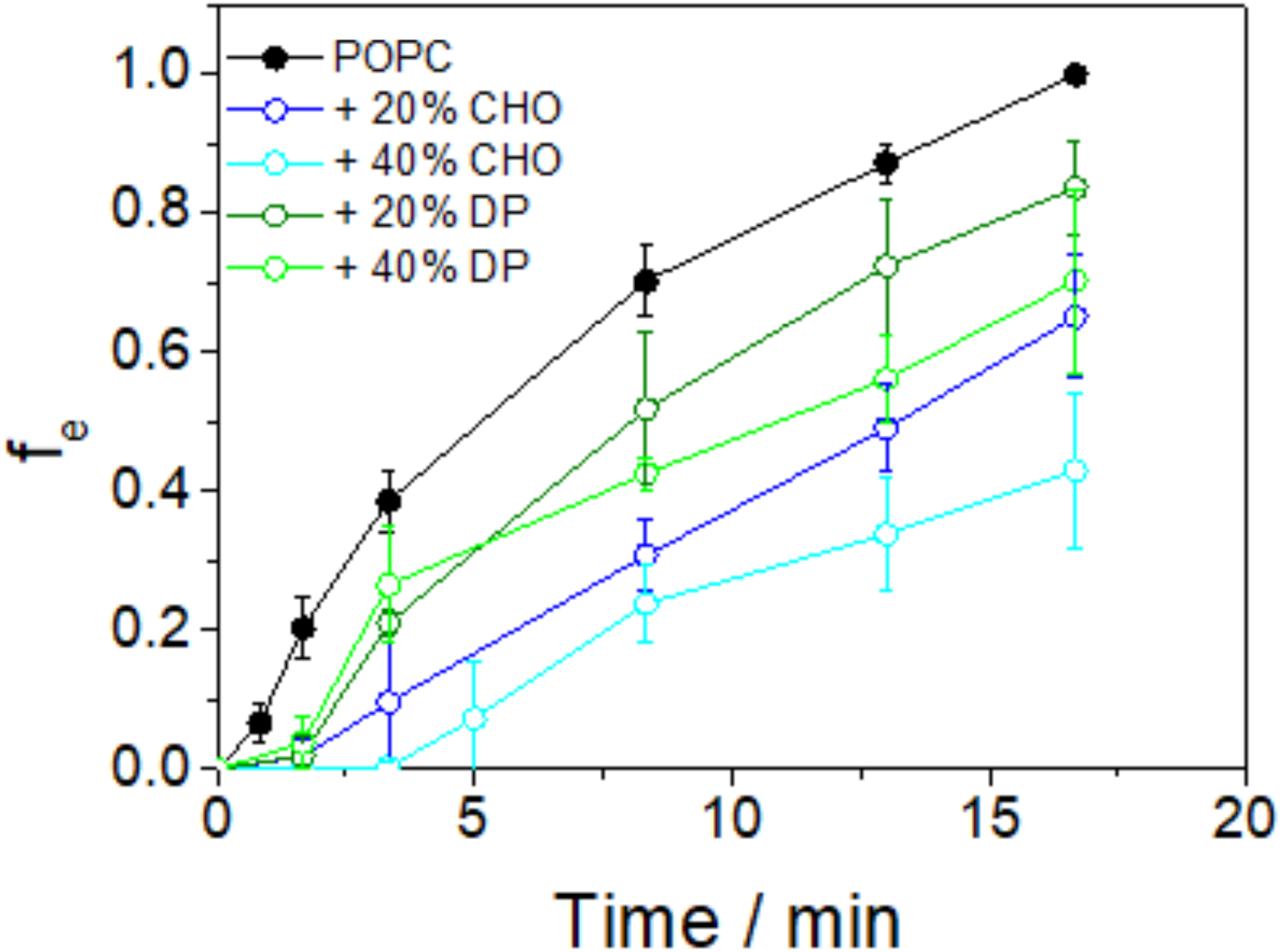
Fraction of GUVs with peptide in their interior as a function of time. All data correspond to the average (± SD) of three independent experiments, using at least 15 GUVs in each experiment.

**Figure SI 9.**
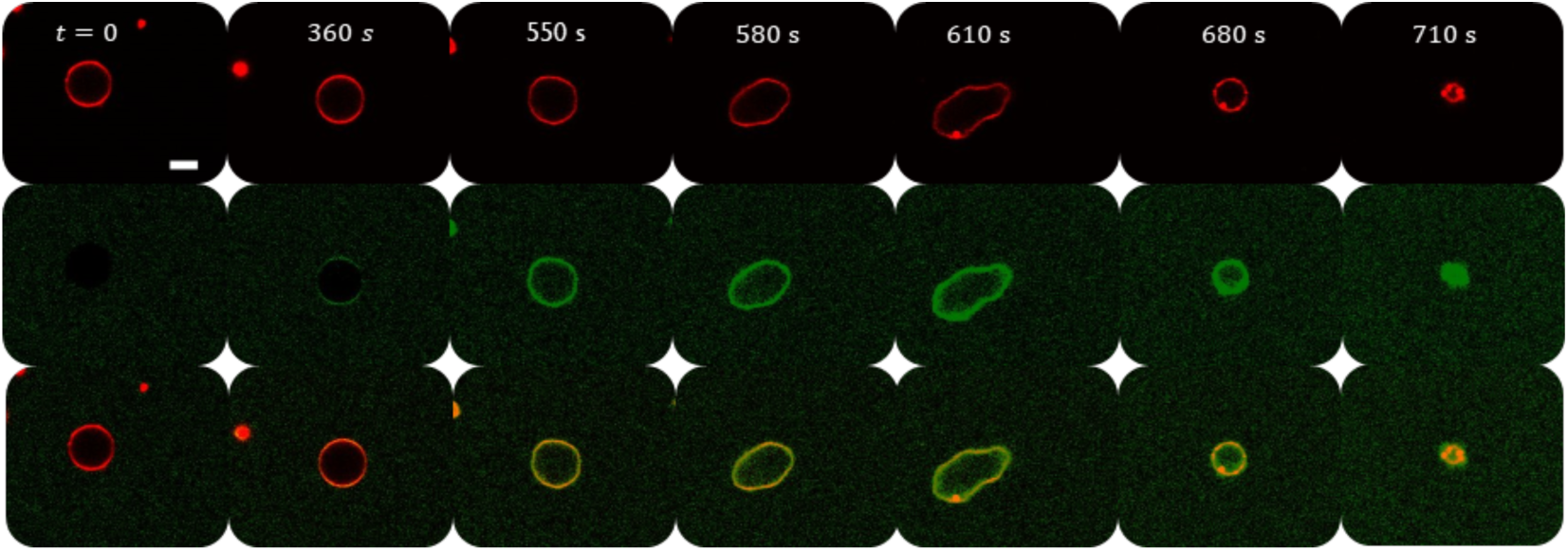
Representative confocal fluorescence images of a GUV composed of POPC doped with 0.5 mol% of PE-Rhodamine (upper panels – red channel) after the addition of MP1 (6 *µ*M) labeled with FITC (middle panels – green channel). The lower panels show the merge of red and green channels. Scale bar: 10 *µ*m. For better visualization of the images, the brightness and contrast range was reduced from an original range of 0–255 to 0–132.

**Figure SI 10.**
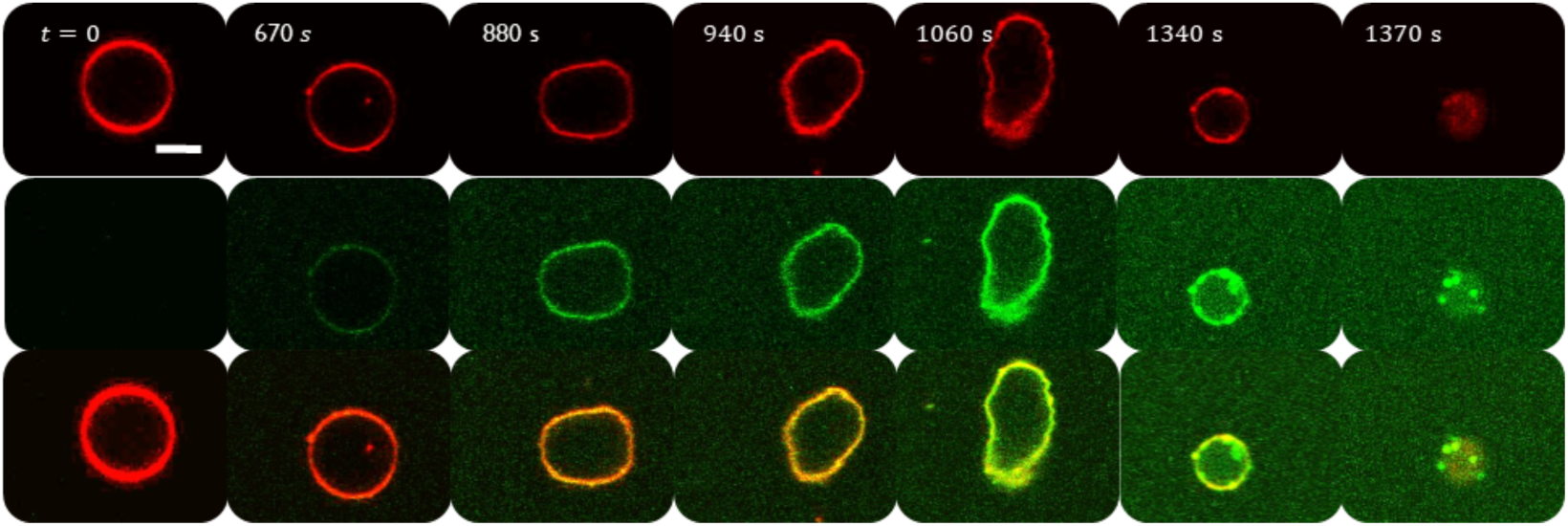
Representative confocal fluorescence images of a GUV composed of POPC + 20% CHO doped with 0.5 mol% of PE-Rhodamine (upper panels – red channel) after the addition of MP1 (6 *µ*M) labeled with FITC (middle panels – green channel). The lower panels show the merge of red and green channels. Scale bar: 10 *µ*m. For better visualization of the images, the brightness and contrast range was reduced from an original range of 0–255 to 0–132.

**Figure SI 11.**
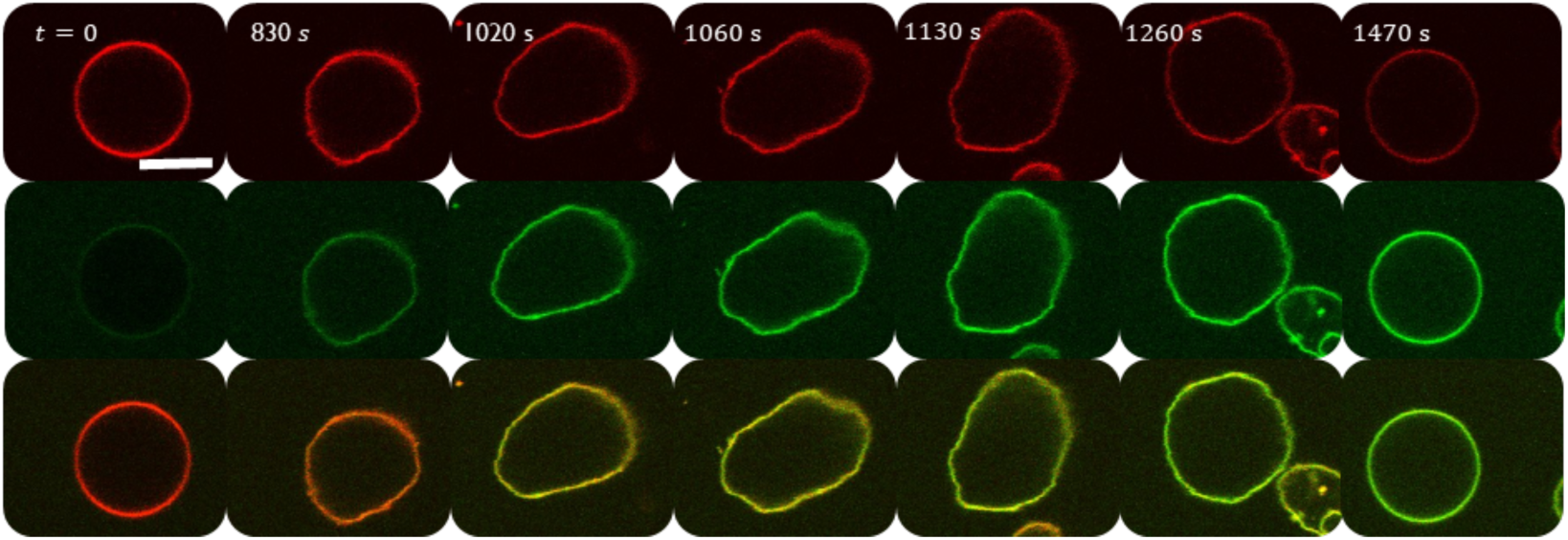
Representative confocal fluorescence images of a GUV composed of POPC + 20% DP doped with 0.5 mol% of PE-Rhodamine (upper panel – red channel) after the addition of MP1 (6.0 *µ*M) labeled with FITC (middle panel – green channel). The lower panels show the merge of red and green channels. Scale bar: 10 *µ*m. For better visualization of the images, the brightness and contrast range was reduced from an original range of 0–255 to 0–132.

**Figure SI 12.**
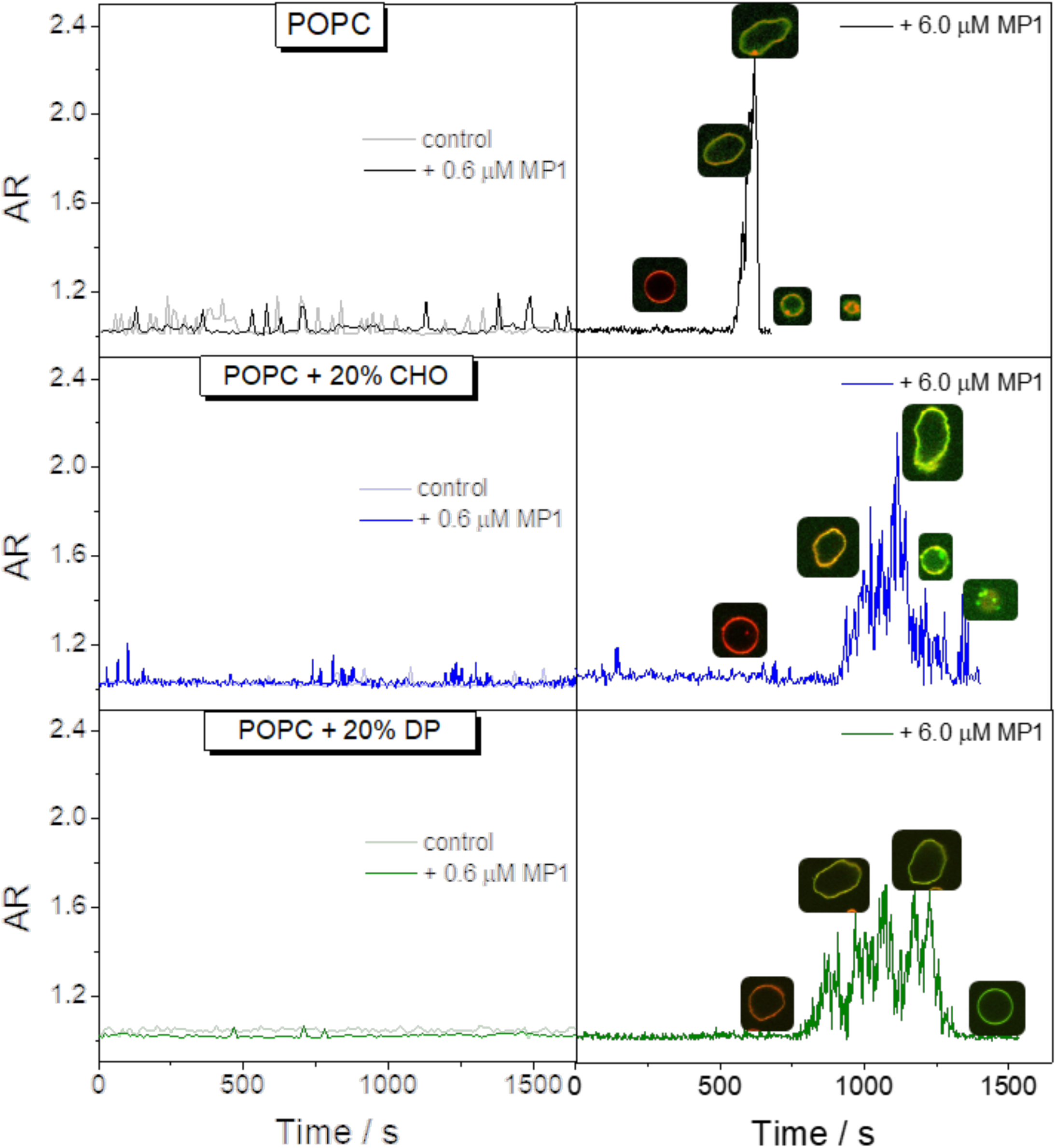
Representative curves showing the aspect ratio (AR) of vesicles as a function of time without peptide or after the addition of 0.6 or 6.0 *µ*M fMP1.

**Figure SI 13.**
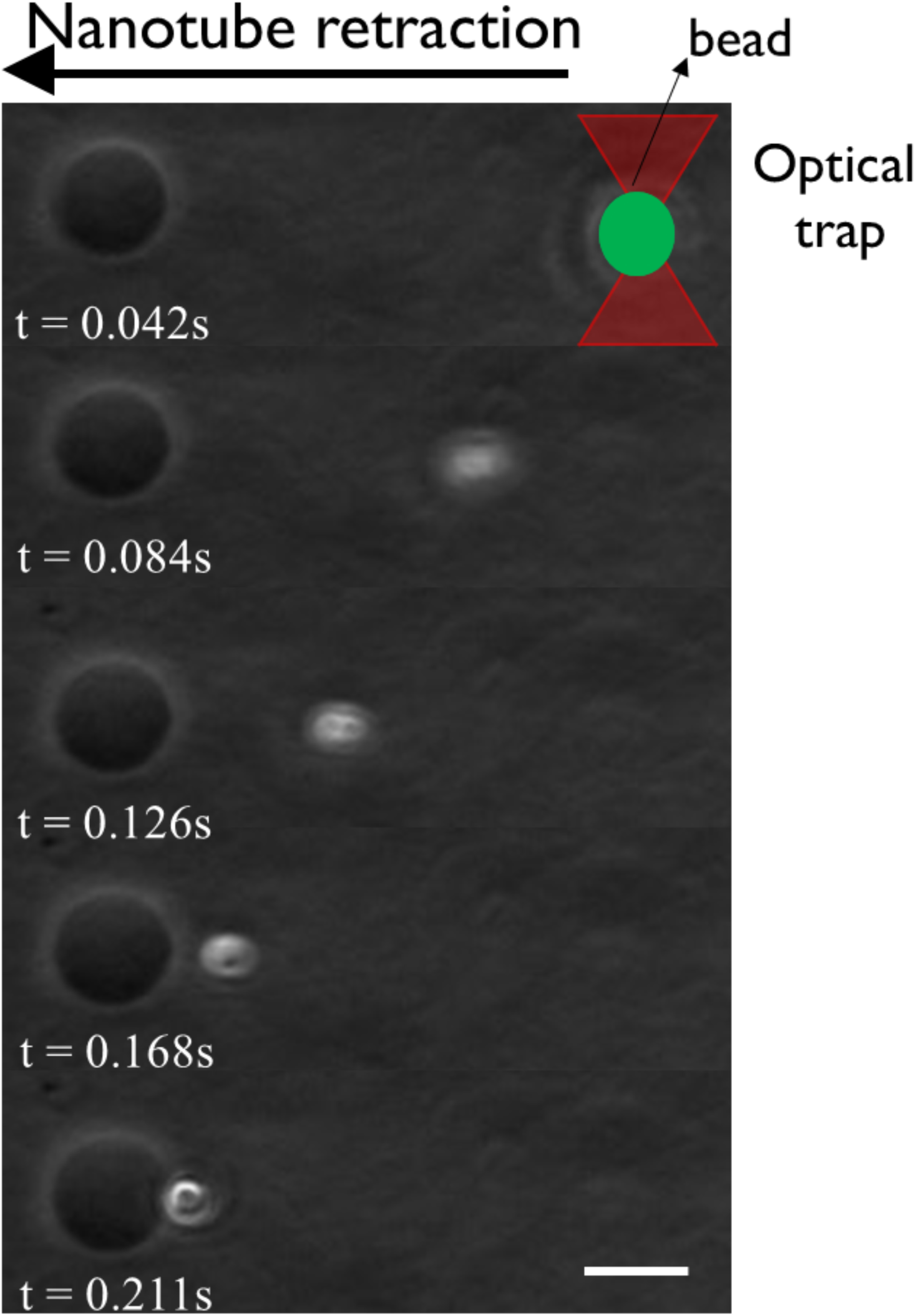
DIC-micrographs showing the kinetic of tether retraction after being pulled from a single biotinylated-POPC GUV immobilized onto an avidin-coated glass slide. Scale bar: 10 *µ*m.

**Figure SI 14.**
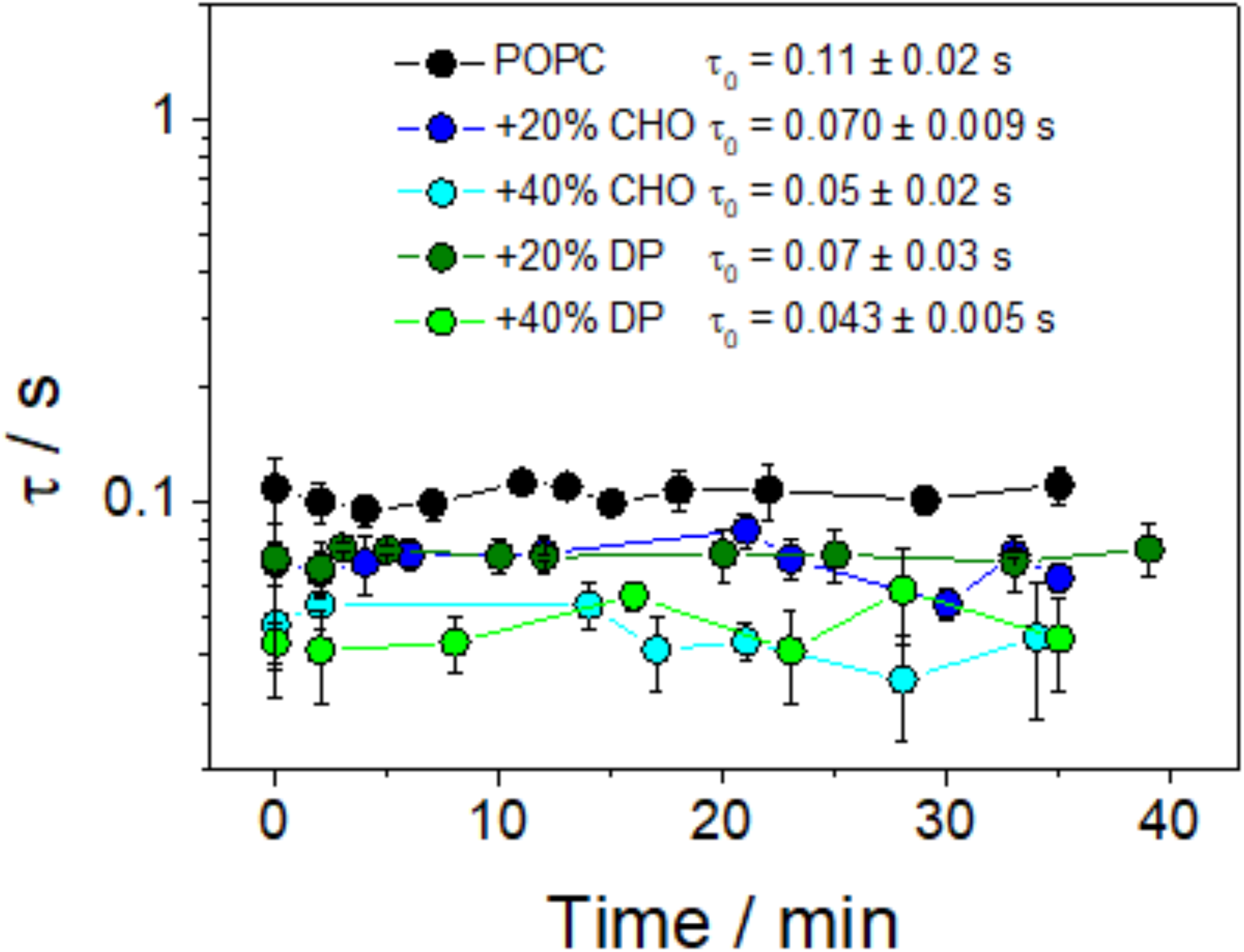
Characteristic times *τ* as a function of time for GUVs composed of pure POPC (black), POPC + 20% CHO (blue), POPC + 20% DP (olive), POPC + 40% CHO (cyan), or POPC + 40% DP (green). Data correspond to the average (± SD) of at least 4 GUVs from at least two independent experiments.

